# An SPFH Protein Couples Membrane Stress to Differentiation in *Bacillus subtilis*

**DOI:** 10.64898/2026.02.06.704340

**Authors:** Sarah S M Baur, Urška Repnik, Tobias Busche, Louisa Rau, Alisa Mondry, Marc Bramkamp

## Abstract

*Bacillus subtilis* adapts to fluctuating environmental stress, such as membrane perturbation or alkaline conditions, using membrane-associated regulatory complexes. Here, we rename the previously termed *pspA-ydjGHI* operon to *pspA-samGHI* (for starvation <u>a</u>nd <u>m</u>otility) to reflect its functional roles in membrane envelope stress signalling. The SamG–SamH membrane proteins recruit SamI, a cytosolic SPFH protein, which stabilizes focal membrane localization and recruitment of PspA, an ESCRT-III homolog. Under normal conditions, this system transiently assembles at the membrane, stabilizing it and allowing proper motility, secretion, and biofilm formation. Loss of SamI (Δ*samI*/Δ*ydjI*) leads to unbalanced SamG–SamH activity leading to a constitutive stress signalling, and global transcriptional changes reminiscent of starvation situations. This, in turn, blocks secretion of the matrix protein BslA, preventing biofilm formation, and reducing motility. Deletion of *samH* in combination with Δ*samI* restores biofilm formation, while Δ*pspA* mutants form biofilms normally, indicating that PspA is dispensable for the developmental phenotype. Our findings reveal that beside membrane integrity SamGHI coordinates transcriptional homeostasis and multicellular development through formation of a membrane integral stress sensor complex.

## 1. Introduction

*Bacillus subtilis* inhabits diverse environments, ranging from soil niches such as the plant rhizosphere to the gastrointestinal tract of animals (Earl *et al*., 2008, Stülke *et al*., 2023, Akinsemolu *et al*., 2024). These habitats are characterized by fluctuating conditions that require rapid physiological adaptation. In response to environmental cues, *B. subtilis* differentiates into distinct phenotypes that allow it to cope with, escape, or endure stress (Lopez & Kolter, 2010, Qin *et al*., 2022). One common lifestyle adopted in nature is biofilm formation, in which cells are embedded in a matrix composed of proteinaceous and carbohydrate-rich fibers which protect the biofilm community from external insults (Vlamakis *et al*., 2013, Mielich-Süss & Lopez, 2015, Arnaouteli *et al*., 2021). Biofilm development is controlled by a complex regulatory network with Spo0A as the central master regulator (Branda *et al*., 2001a, Hamon & Lazazzera, 2001). Activation of membrane-associated sensory kinases triggers a phosphorelay culminating in Spo0A phosphorylation, which in turn inactivates the repressors AbrB and SinR. Their inactivation induces the expression of the matrix gene clusters *epsA–O, tapA–sipW–tasA*, and *bslA* (Hamon *et al*., 2004, Kearns *et al*., 2005, Verhamme *et al*., 2009, Kovacs & Kuipers, 2011). The resulting extracellular polysaccharides and TasA amyloid fibers constitute the core structural components of the biofilm matrix, while BslA forms a hydrophobic surface layer that protects the biofilm (Branda *et al*., 2001b, Branda *et al*., 2006, Kobayashi & Iwano, 2012).

In addition to Spo0A-dependent regulation, alternative sigma factors further fine-tune cellular differentiation. *B. subtilis* encodes 18 alternative sigma factors that modulate transcriptional responses to diverse stresses (reviewed in (Collins *et al*., 2023)). Among them, σ^W^ is activated by membrane stress such as phage infection, antibiotic exposure, and alkaline or high-salt conditions (Petersohn *et al*., 2001, Wiegert *et al*., 2001, Wenzel *et al*., 2012). σ^W^ regulates 37 operons, including three that encode SPFH (Stomatin/Prohibitin/Flotillin/HflK-C) family proteins (Wiegert *et al*., 2001, Cao *et al*., 2002).

SPFH proteins are widespread in all domains of life and are defined by a conserved SPFH domain (Langhorst *et al*., 2005, Rivera-Milla *et al*., 2006, Hinderhofer *et al*., 2009, Bramkamp & Lopez, 2015). Although their precise functions remain incompletely understood, they are broadly associated with membrane organization and homeostasis (Tatsuta *et al*., 2005, Browman *et al*., 2007, Boehm *et al*., 2009). The SPFH domain promotes self-oligomerization (Tavernarakis *et al*., 1999, Boehm *et al*., 2009), leading to the formation of ring-like megacomplexes that associate with cellular membranes (Ma *et al*., 2022, Tan *et al*., 2024).

Two flotillin homologues in *B. subtilis*, FloA and FloT, have been studied in detail. Both possess an N-terminal membrane anchor and a C-terminal flotillin domain following the SPFH region (Bach & Bramkamp, 2015, Bramkamp & Lopez, 2015). FloA and FloT contribute to membrane fluidity, and their absence results in membrane rigidification (Bach & Bramkamp, 2013, Zielinska *et al*., 2020, Álvarez-Mena *et al*., 2025). Loss of these proteins also affects multiple differentiation processes in *B. subtilis*, including sporulation, biofilm development, motility, competence, and protein secretion (Donovan & Bramkamp, 2009, Yepes *et al*., 2012, Dempwolff *et al*., 2012, Bach & Bramkamp, 2013). A third SPFH protein, SamI (formerly YdjI), is encoded at the terminal position of the *pspA*–*samGHI* operon. In this study, we refer to YdjI as SamI (for starvation <u>a</u>nd <u>m</u>otility) based on the observed phenotypes upon deletion and accordingly rename SamG and SamH. SamI differs structurally from FloA and FloT: it contains a C-terminal zinc finger motif and lacks an intrinsic membrane anchor, instead carrying a low-complexity region between the SPFH domain and the zinc finger motif (Savietto Scholz *et al*., 2021). Deletion of *samI* decreases membrane fluidity *in vivo*. Membrane association of SamI is mediated by the integral membrane proteins SamG and SamH. SamI is additionally required for the membrane recruitment and foci formation of PspA (Savietto Scholz *et al*., 2021), an ESCRT-III family protein with roles in membrane maintenance and remodelling (McCullough *et al*., 2015, Thurotte *et al*., 2017). ESCRT-III proteins form large oligomers capable of binding and deforming membranes, supporting membrane fusion and fission, and the stabilization of stressed membrane regions (Heidrich *et al*., 2017, Thurotte *et al*., 2017, Liu *et al*., 2021, Junglas *et al*., 2021, Junglas *et al*., 2025b, Hudina *et al*., 2025, Junglas *et al*., 2025a, Melnikov *et al*., 2025).

Together, the PspA–SamGHI complex is thought to function in membrane stress sensing or response (Savietto Scholz *et al*., 2021, Bramkamp & Scheffers, 2023). The genetic organization and the proposed function of the PspA-SamGHI operon are reminiscent of other phage shock stress response modules found in a variety of prokaryotes. The most studied example is the PspABCF system from *Escherichia coli*. This system is composed of the membrane integral components PspB and PspC which are thought to form a sensory module for detection of cell envelope stress (Kleerebezem *et al*., 1996, Flores-Kim & J., 2016). Upon signal perception PspA oligomerizes into larger complexes, thought to protect membrane integrity (Engl *et al*., 2009). Indeed, deletion of *pspA* has been shown to cause proton leakage and loss of membrane potential (Kleerebezem *et al*., 1996, Kobayashi *et al*., 2007). Despite the similar organization, a direct comparison between the PspBC sensory module and SamGH reveals that there is no protein homology, arguing against referring to the *B. subtilis* proteins as Psp proteins.

Although SamI has been implicated in membrane organization (Savietto Scholz *et al*., 2021), its role in *B. subtilis* developmental programs remains largely unexplored. Here, we investigate SamI’s function during *B. subtilis* cell differentiation. We show that SamI affects multiple developmental processes, including biofilm morphology and cell motility. Our data suggest a function of the SamGHI system in stress sensing and a chronic stress signal leading to a starvation phenotype upon loss of SamI. Importantly, deletion of the entire operon suppresses the observed phenotype, indicating that SamI is required to regulate SamGH signal perception either spatially or temporally.

## 2. Materials and methods

### Strains and media

*Bacillus subtilis* strains derived from the WT168 or the NCIB 3610 background are listed in table S1 and were grown in lysogeny broth (LB medium) (5 g/L yeast extract, 5 g/L NaCl, 10 g/L tryptone) or in MD medium (10.7 g/L K_2_HPO_4_, 6 g/L KH_2_PO_4_, 1 g/L Na_3_C_6_H_5_O_7_ x 5H_2_O, 2 % glucose, 50 μg/mL L-tryptophan, 11 μg/mL ferric ammonium, 2.5 mg/mL L-aspartate, 3 mM MgSO_4_) supplemented with 1 µL/mL casamino acids at 37 °C under shaking conditions. *Escherichia coli* strains (Table S2) were propagated in Lennox broth (5 g/L yeast extract, 10 g/L NaCl, 10 g/L tryptone) at 37 °C under shaking conditions. Growth medium were supplemented with kanamycin (5 µg/mL), erythromycin (1 µg/mL), tetracycline (10 mg/mL), spectinomycin (100 µg/mL), chloramphenicol (5 µg/mL), carbenicillin (100 µg/mL), IPTG (1 mM), or benzyl alcohol (0.05-0.2 %) when necessary. Plates were solidified with 1.5 % agar unless stated otherwise.

### Strain construction

All plasmids generated in this study are listed in table S3 and oligonucleotides used in this study are listed in table S4.

Most single gene deletion strains derived from the *Bacillus subtilis* gene deletion library (Koo *et al*., 2017). These strains were generated by transforming genomic DNA isolated from library strains into the WT168 and NCIB 3610 background using their natural competence of *B. subtilis*. Generation of all knock-out strains of the *pspA-samGHI* operon were verified with primer SB001 and SB052. Flotillin mutants Δ*floT* (Donovan & Bramkamp, 2009), Δ*floA* (Bach & Bramkamp, 2013), and the full operon knock-out Δ*pspA-samGHI* were transformed in the here used WT168 generating SB002 (Δ*floT*), SB071 (Δ*floA*), and SBB007 (Δ*pspA-samGHI*), respectively.

In brief, single colonies were inoculated in 10 mL MD medium supplemented with 50 µL casamino acids (20 %) and grown to an OD_600_ of 1.0-1.5. 10-15 mL MD medium lacking casamino acids were added and after 1 h of incubation 800 µL of the cell culture were transferred and the donor DNA was added to the culture. After 20 min of incubation 25 µL of casamino acids were added and strains were further incubation for 60-80 min before plating on LB plates with respective antibiotics. Transformation plates were incubated ON at 37 °C. If necessary integration into the *amyE* locus was verified with 0.3-1 % starch plates testing for amylases sensitivity and intact *ldh* gene was verified with primer SB137 and SB138 (Dierksheide & Li, 2024).

For removal of the antibiotic cassette strains generated with the gene deletion library were transformed with the plasmid pDR244 following the description of (Koo *et al*., 2017). Loss of antibiotic cassette was confirmed by sequencing.

To generate genomic DNA a fresh colony from the respective strain was inoculate in 2.5 mL of LB medium and grown until the culture lost its motility for 3-4 h (adapted from (Ward & Zahler, 1973)). Cell pellets were resuspended in 1 mL SCC (0.15 M NaCl, 0.01 M Na_3_C_6_H_5_O_7_, pH 7.0) supplemented with 0.2 mg/mL lysozyme and incubated for 15 min at 37 °C until cell suspension was lysed. 1 mL of 4 M NaCl were added to induce osmolysis. The lysed cell suspension was filtered (0.45 μm) and stored at −20 °C if necessary.

The plasmids used in this study (Table S3) were generated with Gibson Assembly (Gibson *et al*., 2009) with the NEBuilder® HiFi DNA Assembly Master Mix (E2621). Fragments for Gibson Assembly were amplified using Phusion High-Fidelity DNA Polymerase (M0350) or Q5 High-Fidelity DNA Polymerase (M0491). Plasmids based on pKill were verified with primers SB118 and SB119 and plasmids based on pDR111 were verified with SB035 and SB036.

For double and triple deletions of the *pspA-samGHI* operon genes the primers SB159 to SB170 were used. gDNA from SBB001 and SBB070 and plasmids pKill-*samI-halo-spec* (SBE023) and pKill-*spec-halo-samI* (SBE024) were used as template. Primers SB165 to SB170 were used for pKill-samHI, primers SB159 to SB164 were used for pKill-*samGH*, and primers SB160 to SB166 were used for pKill-*samGHI*. The resulting plasmids were transformed into competent *E. coli* DH5α and plasmids were isolated, verified, and sequenced from the resulting colonies. Up on verification of the correct sequence, plasmids were linearized and transformed into SBB001, generating SBB138 (Δ*samHI*), SBB159 (Δ*samGH*), and SBB160 (Δ*samGHI*).

For complementation with *samI* under an IPTG inducible promoter in the *amyE* locus the primers SB027 to SB030 were used and the plasmid pDRyuaB2 (Kovacs & Kuipers, 2011) and genomic DNA of SBB001 was used as a template. The resulting plasmid pDR111-*samI* was transformed into competent *E. coli* DH5α and from the resulting and verified colonies plasmids were isolated and sequenced. Up on verification of the correct sequence, plasmids were linearized and transformed into SBB003, generating SBB060 (Δ*samI* P_hs_-*samI*). Transformation of genomic DNA from SBB070 into SBB060 resulted in SBB067 (Δ*samG* Δ*samI* P_hs_-*samI*).

SamH was complemented via an *amyE* integration of *samH* under an IPTG inducible promoter. Primers SB028 and SB257 to SB259 were used and the plasmid pDRyuaB2 (Kovacs & Kuipers, 2011) and genomic DNA of WT168 were used as a template. The resulting plasmids pDR111-*samH* was transformed into competent *E. coli* DH5α and from the resulting colonies plasmids were isolated and sequenced. Upon verification of the correct sequence, plasmids were linearized and transformed into SBB003, generating SBB193 (Δ*samI* P_hs_-*samH*).

To generate the fragments for BslA-His expression (*bslA::bslA-6xHis-cat*) primers SB037 to SB038, SB104 to SB106, SB111, SB114 to SB115, SB118, and SB119 were used. Fragments generated with primers SB037 to SB038, SB104 to SB106, SB111, SB114 to SB115 were used for Gibson assembly and the resulting construct was amplified again with SB118 and SB119. This fragment was verified by sequencing and directly transformed into *B. subtilis* SBB041 generating SBB131. DNA was isolated from SBB131 and retransformed into SBB001 and SBB003 generating SBB127 (BslA-His) and SBB129 (Δ*samI* BslA-His), respectively.

For generation of a DivIVA-GFP fusion DNA was isolated from 1803 (Edwards *et al*., 2000) and transformed into SBB001 and SBB003 generating SBB133 (DivIVA-GFP) and SBB134 (Δ*samI* DivIVA-GFP).

For the *fliM*-GFP fusions gDNA of a Δ*fliM* from the gene deletion library was transformed into SBB001 and SBB094, generation SBB149 (Δ*fliM*) and SBB150 (Δ*samI* Δ*fliM*). These strains were further transformed with gDNA isolated from the strain DS8521 (Guttenplan *et al*., 2013) and generating SBB151 (Δ*fliM* P_fla/che_ FliM-GFP) and SBB152 (Δ*samI* Δ*fliM* P_fla/che_ FliM-GFP), respectively.

### Growth curves

Overnight cultures were inoculated from a fresh plate in MD or LB medium with respective antibiotics. Cultures were diluted to an OD_600_ of 0.1 in the next morning and growth was monitored over the next 8 h. An additional timepoint was measured after 24 h.

### Biofilm assay

The plates for biofilm assays were poured in the morning before inoculation consisting of LBGM medium (LB medium supplemented with 1mM MnSO_4_ and 1 % (v/v) glycerol (Shemesh & Chai, 2013)) or MSgg medium (1.925 mM KH_2_PO_4_ (pH 7.0), 3.075 mM K_2_HPO_4_ (pH 7.0), 100 mM MOPS (pH 7.0), 0.5 % glutamate, 2 mM MgCl_2_, 700 µM CaCl_2_, 50 µM MnCl_2_, 1 µM ZnCl_2_, 2 µM thiamine, 50 µg/mL phenylalanine, 50 µg/mL tryptophan, and 0.5 % glycerol). 50 µM FeCl_3_ were added to MSgg medium on the day of the experiment (Branda *et al*., 2001b). Cultures were grown by picking a fresh colony and inoculating it into LB medium. After reaching an OD_600_ of 0.4 to 0.8 cultures were diluted to an OD_600_ of 0.1, IPTG (1 mM) were added when necessary, and grown until they reached an OD_600_ of 0.8 to 1.0 again. The OD_600_ was normalized to 0.8 and 1.5 µL were inoculated onto LBGM or MSgg plates or into 1.5 mL LBGM medium in a 24-Well plate. Agar and medium were supplemented with 0.05-2 % benzyl alcohol or 1 mM IPTG when indicated. The biofilm plates were incubated at 30 °C for 48 h (LBGM based medium) or 72 h (MSgg based medium) and the pellicle development was observed at RT for one week. Growth and GFP fluorescence were documented with a ChemiDoc™ MD. For further isolation of RNA or protein biofilms were harvested after 48 h.

Contact angles of biofilms were determined after 48 h (for LBGM) or 72 h (for MSgg) of growth by pipetting a 5 µL drop of deionized water, stained with Coomassie Brilliant Blue for increased contrast. The drop was left for 5 min to equilibrate before an image was taken from the side with a camera of a Pixel 7a. Contact angles were analyzed with the ImageJ plugin Contact Angle.

### Motility assays

Motility assays were performed on LB plates with varying agar concentrations. 0.5 % agar induced swimming behavior, whereas 0.7 % agar resulted in swarming behavior. Plates were poured the day before and dried for 30 min. 20 min before inoculation plates were dried again.

Overnight cultures were inoculated from a fresh plate into LB medium with respective antibiotics if necessary. The cultures were diluted 1:100 the next morning and grown until an OD_600_ of 0.5-0.8. Cultures were diluted to an OD_600_ of 0.1 and after reaching an OD_600_ of 0.5-0.8 again, 5 µL of the cultures were inoculated onto the center of the LB plates. Plates were dried for 10 min and placed at 37 °C. Plates from swimming assays were kept in a humidity chamber, an airtight sealed box with a reservoir of water in it. Development was observed for up to 48 h and documented with a ChemiDoc™ MD.

### Fluorescence Microscopy

For determination of FliM-GFP expression cells were picked from the center and periphery of the biofilm and resuspended in 5 µL PBS. 2 µL of cell suspension were added on a 1 % agarose pad in PBS and imaged at the Axio Zeiss Imager M1 fluorescence microscope (EC Plan-Neofluor 100x/ 1.30 Oil Ph3 objective) equipped with an AxioCam HRm camera. GFP was excited at 470 nm for 500 ms and fluorescence emission was captured at 509 nm. Digital images were acquired with AxioVision Release 4.6.3 and analyzed and edited with Fiji ImageJ.

### Electron Microscopy

Fixative was poured onto a biofilm growing on LBGM agar. For ultrastructural analysis by TEM or by SEM, samples were fixed with 1 % glutaraldehyde (GA) in 200 mM HEPES (pH 7.4) for several days. After 10 min of initial fixation, biofilms were cut into narrow strips to improve fixation efficiency.

For epoxy resin embedding, samples were postfixed with 1 % osmium tetroxide prepared in 1.5 % potassium ferricyanide for 1.5 h on ice, stained with 2 % uranyl acetate for 2 h and dehydrated using a graded ethanol series (50-70-80-90-96-100 %) and then acetone, all steps at room temperature for a minimum of 15 min per step. Samples were progressively infiltrated with epoxy resin over two days before resin polymerization. Ultrathin 80-nm sections were cut using a Leica UC7 ultramicrotome and a Diatome ultra diamond knife and were contrasted with saturated aqueous uranyl acetate solution and Reynolds’ lead citrate. Grids were imaged with a Tecnai G2 Spirit BioTWIN transmission electron microscope (FEI, now Thermo Fisher Scientific), equipped with a LaB6 cathode and operated at 80 kV using the TEM User Interface software (v. 4.2 build 6173). Images were acquired with a CCD side-entry MegaView III G2 camera using iTEM software (v. 5.0) (all Olympus Soft Imaging Solutions, now EMSIS). To prepare 1 µm semi-thin epoxy resin sections, few alterations were made to the above protocol. Specifically, 0.015 % ruthenium red and 50 mM L-Lysine were added to the fixative for the initial 30 min fixation, and then fixation was continued in the fixative supplemented with only 0.015 % ruthenium red. Osmium tetroxide postfixation was also carried out in the presence of 0.075 % ruthenium red. Sections were mounted on microscope slides and stained with Richardson’s solution (alkaline solution of azure II (50%) and methylene blue (50%)) (Richardson *et al*., 1960). Bright-field images were acquired using a Zeiss Primostar microscope equipped with a Plan-Achromat 40x objective (0.65 N.A.) and an Axiocam 105 color camera, which was operated with a Zen Blue (v. 3.2.) software.

For SEM analysis, samples were first dehydrated with a graded ethanol series and then subjected to critical point drying using a CPD 030 (Bal-tec). Dried specimens were mounted on adhesive carbon tape and sputter coated with a 3 nm platinum layer using a Safematic CCU 010 HV compact coating unit. Samples were imaged with a Zeiss Sigma 300 VP SEM operated at 1 kV accelerating voltage and using a secondary electron detector.

### RNA Isolation

Biofilms were transferred into a tube and submerged in 800 µL DNA/RNA Shield supplemented with 10 mg/mL lysozyme. To homogenize biofilms were first passed through a 23g1 needle 10-15 times and then sonicated for 12 bursts (duty cycle 20 %, output control 2). 30 min incubation at 37 °C under shaking conditions and subsequent sonification for 4×20 bursts lysed the cell, and debris were pelleted by centrifugation at max. speed for 2 min. To isolate total RNA the protocol of the *Quick*-RNA™ Miniprep Plus Kit (Zymo Research) was followed. Concentration of isolated RNA was measured with a NanoDrop and integrity was confirmed in a denaturing agarose gel (Aranda *et al*., 2012).

### RT-qPCR

Reverse transcription qPCR was performed using the Luna® Universal One-Step RT-qPCR Kit (NEB E3005) following manufacturers description. 100 ng of RNA were used as template and oligonucleotides used are listed in table 2 (primers SB080 to SB103, SB116 and SB117). Amplification was carried out with AriaMx Real-Time PCR System (Agilent Technologies). *divIVA* amplification was used as reference and the fold change in expression of *spo0A, abrB, lip, degQ, sinR, bslA, epsA, tasA*, and *samI* was determined by 2^−*Δ ΔCq*^.

### RNA preparation

Sample purity and integrity was verified with Xpose and Agilent Bioanalyzer and DNA contaminated samples were treated with DNase Kit (Qiagen) and rechecked. RNA with an RNA Integrity Number (RIN) >9 and an rRNA Ratio [23S/16S] >1.5 and free of DNA was used for sequencing library preparation and sequencing.

### Whole Transcriptome Sequencing

High-quality total RNA aliquots of the biological replicates were rRNA-depleted using Pan-Bacteria riboPOOL (siTOOLs BIOTECH). Successful rRNA depletion was confirmed using Agilent RNA Pico 6000 kit on Agilent 2100 Bioanalyzer (Agilent Technologies). The TruSeq Stranded mRNA Library Prep Kit from Illumina, starting with fragment, prime, and finish was used to prepare the cDNA libraries. The cDNA libraries were then sequenced in paired-end mode on an Illumina NextSeq2000 system with 50 nt read length and a minimum of 5 million cluster per sample.

The resulting sequence reads were trimmed with Trimmomatic v0.33 (Bolger *et al*., 2014) to a minimal length of 35 base pairs and subsequently mapped onto the *Bacillus subtilis* subsp. subtilis str. 168 genome reference sequences (NC_000964.3) using Bowtie 2 (Langmead & Salzberg, 2012). The ReadXplorer software version 2.0 (Hilker *et al*., 2016) and the integrated DESeq2 algorithm (Love *et al*., 2014) were used for the visualization of the mapped reads and the differential gene expression analysis, respectively. Differentially expressed targets were filtered with a log2 fold change (m-value) ≥ | 1| and an adjusted p-value ≤ 0.01.

### Isolation of biofilm matrix

To isolate matrix proteins from biofilms the following protocol was adjusted (Arnaouteli *et al*., 2017). Three biofilms were combined and resuspended in 800 µl PBS supplemented with protease inhibitor cocktail and DNase I and passaged through a 23g1 needle 10-15 times. Samples were sonicated (duty cycle 20 %, output control 2) for 20 impulses 4-6 times and matrix disruption was verified under a microscope. After centrifugation for 20 min at 9000xg and 4 °C supernatant was filtered (0.2 µm) and contained biofilm matrix while the pellet contained cells. Pellet was resuspended in same volume as before and sonicated by 2×20 impulses (duty cycle 20 %, output control 2). Concentrations were determined with NanoDrop.

### Immunoblotting

100 µg samples were mixed with 4x SDS Loading dye and 10x Laemmli buffer (250 mM Tris, 1.92 M glycine, 1 % SDS, pH 8.7) and incubated for 10 min at 90 °C, before loading into a 14 % SDS PAGE containing 0.6 % TCE. SDS PAGE were run at 120 V and protein content was visualized in a ChemiDoc™ MD. Proteins were transferred to a PVDF membrane (Bio-Rad) activated with absolute ethanol. The membrane was incubated with Penta-His antibody (1:1000, ID: 34660, Qiagen) after blocking and further incubated with goat anti-mouse secondary antibody HRP conjugate (1:5000, ID: 62-6520, Invitrogen) before developing with Pierce™ ECL Western Blotting Substrate (Cat. No. 32109, Thermo Scientific™). Bioluminescence was detected with signal accumulation mode in ChemiDoc™ MD (Bio-Rad).

Quantification of band intensities was performed with ImageJ gel analyser and normalized to band intensity of the WT168. Significances were calculated using one-sample t-test with p-value ≤ 0.05 and results were depicted within the graphs (ns – non-significant, * – p-value ≤ 0.05, and ** – p-value ≤ 0.005).

### Statistical analysis

Origin 2024b (Academic) was used to generate all graphs and conduct data analysis and statistical tests determining significance. Detailed description of statistical testing can be found within the methods.

## 3. Results

### SamI impacts biofilm development

The flotillin-like proteins FloA and FloT are known to influence several developmental processes in *B. subtilis*. Loss of FloT impairs sporulation (Donovan & Bramkamp, 2009, Yepes *et al*., 2012) and motility, while additional deletion of FloA further reduces motility (Dempwolff *et al*., 2012), decreases protein secretion (Bach & Bramkamp, 2013), and diminishes extracellular matrix production during biofilm formation (Yepes *et al*., 2012, Bach & Bramkamp, 2013). In contrast, the role of the third SPFH protein, SamI, in *B. subtilis* development has remained unclear so far. To better understand SamI’s cellular function, we investigated its impact on biofilm formation.

We first analyzed the morphology of Δ*samI* biofilms grown on solid medium. Compared to the wild type strain (WT168), Δ*samI* colonies exhibited reduced structural complexity and smoother edges (Figure 1A). When biofilm formation at the liquid–air interface was tested, Δ*samI* was unable to form a surface-covering pellicle (Figure 1A). A hallmark of the *B. subtilis* biofilm is its highly hydrophobic surface (Arnaouteli *et al*., 2016). Therefore, we quantified surface hydrophobicity by placing a water droplet on the biofilm and measured the contact angle. Wild type cells have hydrophobic colony surfaces with a contact angle of 125.7 ± 6.58° (Figure 1B, Table S5). Δ*samI* biofilms lacked the characteristic hydrophobicity of wild type biofilms. Consequently, the contact angle dropped to zero (Figure 1B, Table S5). This phenotype was independent of the medium used, as similar effects were observed on different biofilm-inducing media (Figure S1, Table S5). Importantly, growth of Δ*samI* in shaking liquid cultures was indistinguishable from the wild type in both minimal and rich media (Figure S2), indicating that SamI’s role becomes relevant only upon initiation of biofilm differentiation.

**Figure 1:**
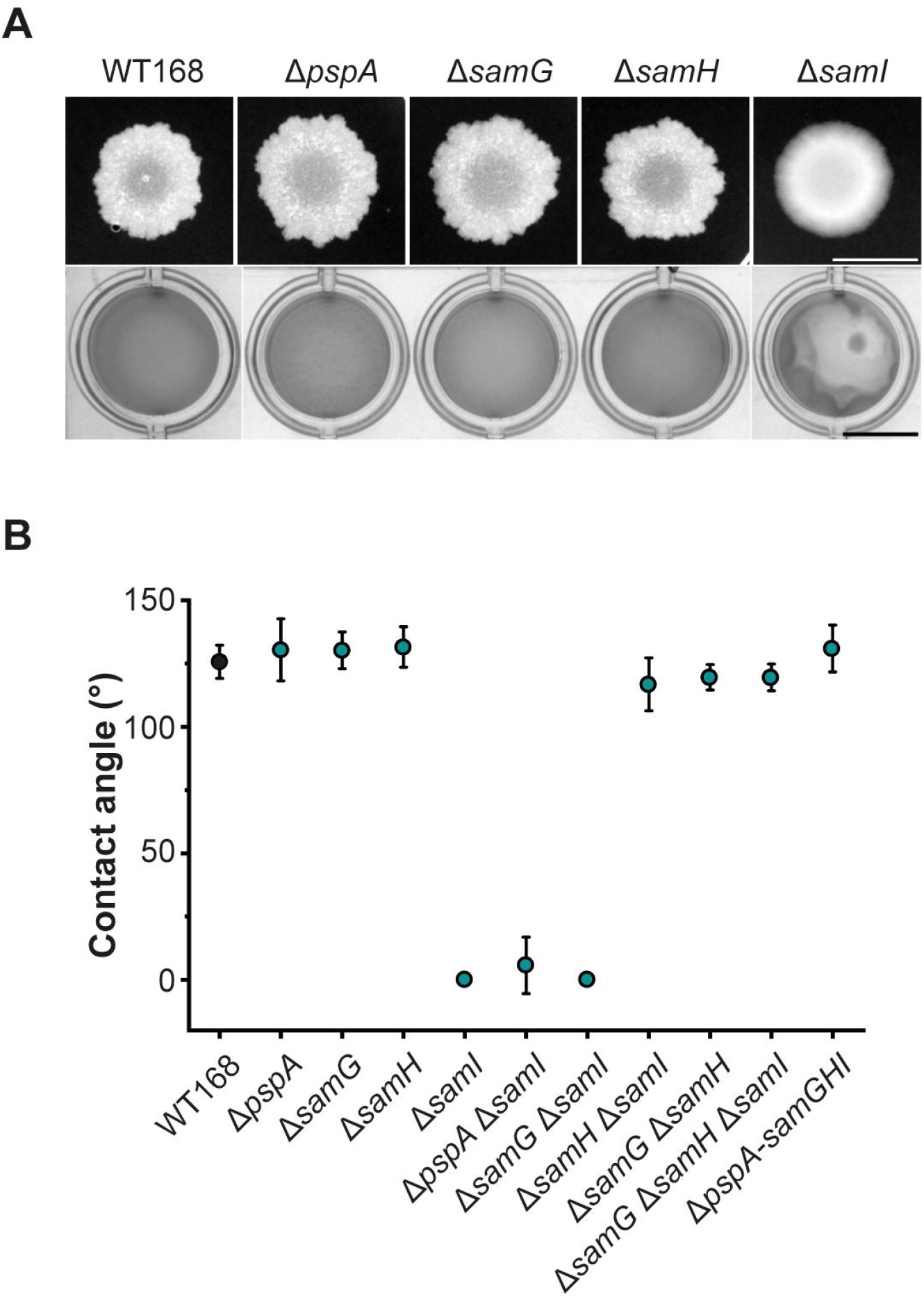
Deficient biofilm development of a *samI mutant on liquid and solid medium*. (A) Biofilm (upper panel) and pellicle growth (lower panel) of WT168 and single mutants of the *pspA-samGHI* operon on LBGM medium (scale: 1 cm). (B) Hydrophobicity measurement of corresponding biofilms and additional double and triple deletions, as well as a full operon knock-out (average and standard derivation was determined for three independent replicates).

SamI is encoded in the *pspA*–*samGHI* operon together with PspA, SamG, and SamH. Because proteins of this operon have been reported to interact closely (Savietto Scholz *et al*., 2021), we additionally tested combinations of deletions. All single mutants, except the deletion of *samI*, revealed wild type like hydrophobicity, colony morphology, and pellicle development (Figure 1A and B, Table S5). Further, a double deletion of *samG* and *samH* also showed no phenotype. The hydrophilic phenotype of Δ*samI* was retained upon additional deletion of *pspA* or *samG* besides *samI*, whereas additional deletion of *samH* in the Δ*samI* background restored hydrophobicity, indicating that the Δ*samI* phenotype depends on the presence of SamH. Consistently, a triple deletion of *samG, samH*, and *samI*, as well as a complete operon knock-out, both produced hydrophobic biofilms. These results suggest that the membrane protein SamH plays a major role in phenotype production, regulated by SamI’s function during biofilm development.

To further explore the loss of biofilm hydrophobicity, we complemented the Δ*samI* strain by expressing *samI* or *samH* ectopically under an inducible promoter. Overexpression of either gene alone failed to restore hydrophobicity (Figure S3A, Table S5). Only when *samG* was additionally deleted, thus, removing a membrane anchor for SamI, induction of *samI* lead to an increase in hydrophobicity (Figure S3A). Since deletion of *samG* did not alter *samI* transcript levels (Figure S3B), this suggests that SamI functions as a regulator of the SamGH complex, thus leading to uncontrolled activity of the SamGH complex in absence of SamI during biofilm development. Together with the genetic interaction data, these findings indicate that absence of SamI could lead to unregulated activity of the membrane components SamGH.

We described before that deletion of *samI* gives rise to decreased membrane fluidity (Savietto Scholz *et al*., 2021). To test the influence of membrane fluidity on biofilm development, we also tested the effect of low concentrations of benzyl alcohol, a membrane-fluidizing agent known to counteract membrane rigidification caused by loss of SPFH proteins (Bach & Bramkamp, 2013, Zielinska *et al*., 2020, Álvarez-Mena *et al*., 2025). In the wild type and in *floA* and *floT* mutants, 0.05 to 0.1 % benzyl alcohol promoted biofilm formation (Figure S4, Table S5). This effect was not observed in the Δ*samI* strain, indicating that SamI’s role in biofilm formation is not related to the membrane-fluidity regulation typically associated with other SPFH proteins.

To gain deeper insight into the morphological differences between wild type and Δ*samI* biofilms, we performed microscopic analyses. Brightfield images of resin sections of wild type biofilm, stained with Richardson solution, revealed a highly structured surface architecture (Figure 2A and S5). An unstained layer of variable thickness was visible at the bottom of the biofilm after staining with Richardson solution (Figure 2A) corresponding to a layer of empty cell envelopes (Figure S5A). In contrast, Δ*samI* biofilms exhibited a uniform, monotonous surface structure and a stable thickness of the empty-cell layer across the cross-section. Higher-resolution transmission electron micrographs of the biofilm cross-sections revealed a thin, electron-dense coat covering only the wild type biofilm which was absent in the Δ*samI* strain (Figure 2B). Scanning electron microscopy confirmed the presence and absence of this surface layer in wild type and *samI* mutant biofilms, respectively (Figure 2C). *B. subtilis* biofilms are known to be coated by a proteinaceous layer formed by BslA, which confers hydrophobic properties to the biofilm (Kobayashi & Iwano, 2012, Hobley *et al*., 2013). Analysis of a Δ*bslA* biofilm revealed that Δ*bslA*, like Δ*samI*, exhibits a loss of architectural complexity (Figure 2A) and lacks the surface coat (Figure 2B), suggesting that the lack of this layer in Δ*samI* likely explains the observed loss of hydrophobicity.

**Figure 2:**
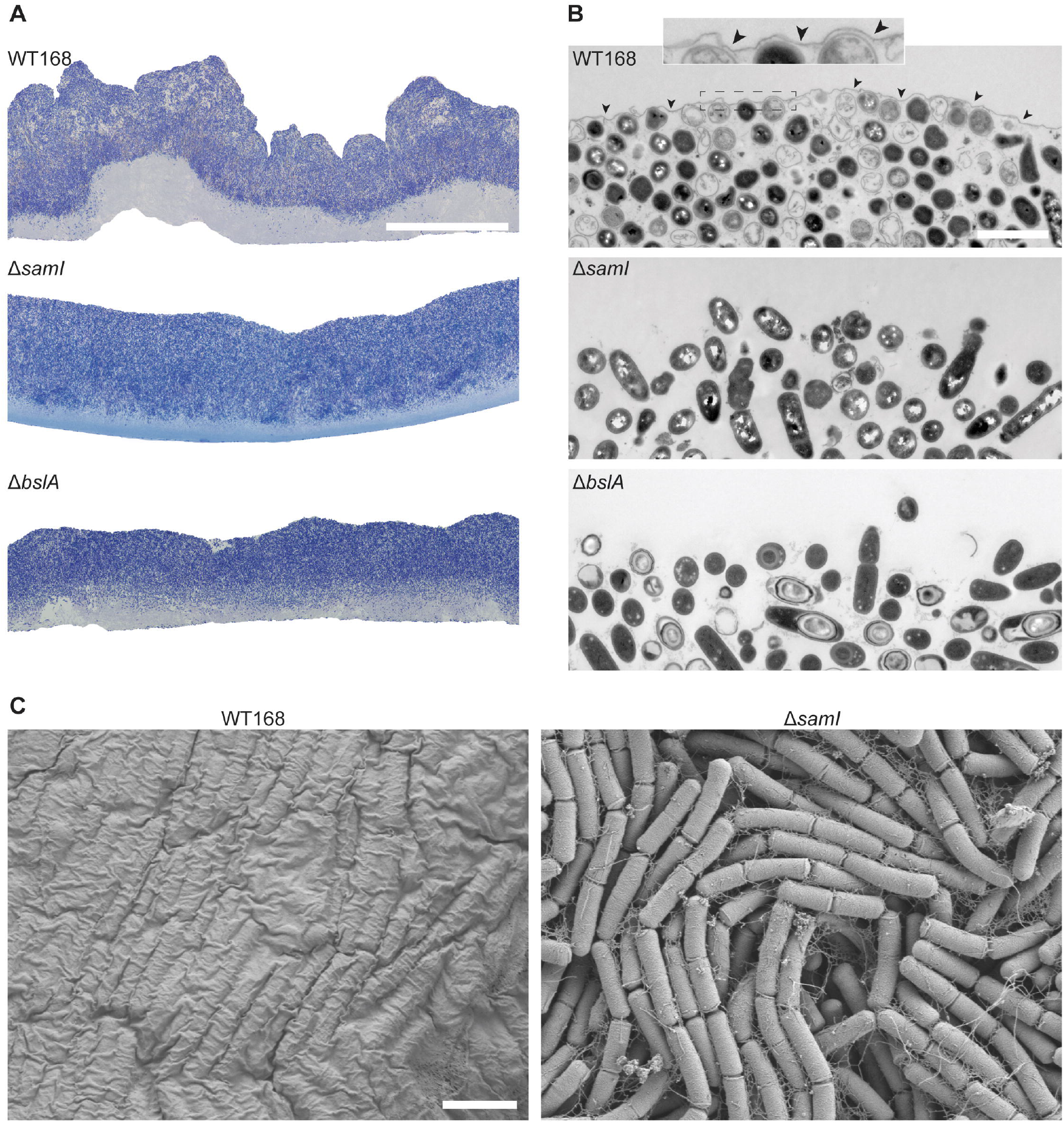
Δ*samI* leads to structural changes in biofilms. (A) Cross-section morphology of WT168, Δ*samI*, and Δ*bslA* biofilms (scale: 100 µm). Brightfield images of 1 µm resin sections stained with Richardson solution. (B) TEM images of the biofilm surface in perpendicular 80 nm resin sections. The surface layer is present in WT168 (arrowheads) but missing in Δ*samI* and Δ*bslA* (scale: 2 µm). (C) SEM top-view images of the biofilm surface in WT168 and Δ*samI*. A continuous surface layer covers bacteria in WT168 (scale: 2 µm).

So far, our results show that SamI is required for proper biofilm development in *B. subtilis*, contributing to both architectural complexity and formation of the hydrophobic BslA layer at the biofilm surface. This function depends on the membrane-anchored proteins SamGH that seem to be regulated by SamI.

### Absence of SamI leads to a starvation phenotype

To understand the effect that SamI has during biofilm development we performed transcriptome analysis, comparing transcript levels between the wild type and Δ*samI* biofilms. In total 4,242 transcripts were detected. Of these transcripts, 62 and 111 (1.5 % and 2.6 %) had significantly increased or decreased levels, respectively (Figure 3A, Table S6). Surprisingly, the expression of *bslA* was upregulated, while the expression of the transition state regulator *abrB* was reduced. A lower expression of the ICEBs1 genes was observed as well. ICEBs1 is an integrative and conjugative genetic element which is activated upon high cell densities and low nutrient availability or global DNA damage, leading to an induction of the SOS response (Auchtung *et al*., 2005). Further a reduction of genes involved in sporulation, motility and chemotaxis, including a strong reduction of the flagellin *hag*, was observed. Additionally, a significant change in expression of gene clusters associated with metabolism was observed, showing an upregulation of the energy-generating part of glycolysis (*tpiA, gapA, pgk, pgm, eno*, and *pdhABCD*) and a corresponding downregulation of the gluconeogenic gene *gapB*. Furthermore, an increase in gene clusters involved in nitrogen and phosphate starvation were observed, including the *narGHIJ* operon encoding a nitrate reductase complex and its chaperone. Several operons associated with phosphate limitation such as: both major alkaline phosphatases (*phoA* and *phoB*) and two phosphodiesterases (*phoD* and *glpQ*) also showed increased expression. All of these genes encode secreted enzymes involved in phosphate scavenging and cell wall teichoic acid turnover. In addition, an induction of the *tuaBCDEFG* operon was detected, encoding a complex responsible for the exchange of teichoic acids with teichuronic acids, lacking the phosphate moieties, and an induction of *tatAD-CD*, encoding the twin arginine translocase, which secretes PhoD. A downregulation of genes associated with the methionine salvage pathway was detected as well. This includes the *mtnKA* operon, the *mtnWXBD* operon, *mtnE, mtnU*, and *speD*, all involved in the recycling of S-adenosylmethionine into methionine. Beside these changes in metabolism associated genes we also observed an increase in *hpf* transcripts. *hpf* encodes for the ribosome hibernation factor that inactivates ribosomes to reduce translation under stress conditions (Drzewiecki *et al*., 1998, Tagami *et al*., 2012, Akanuma *et al*., 2016, Feaga *et al*., 2020).

**Figure 3:**
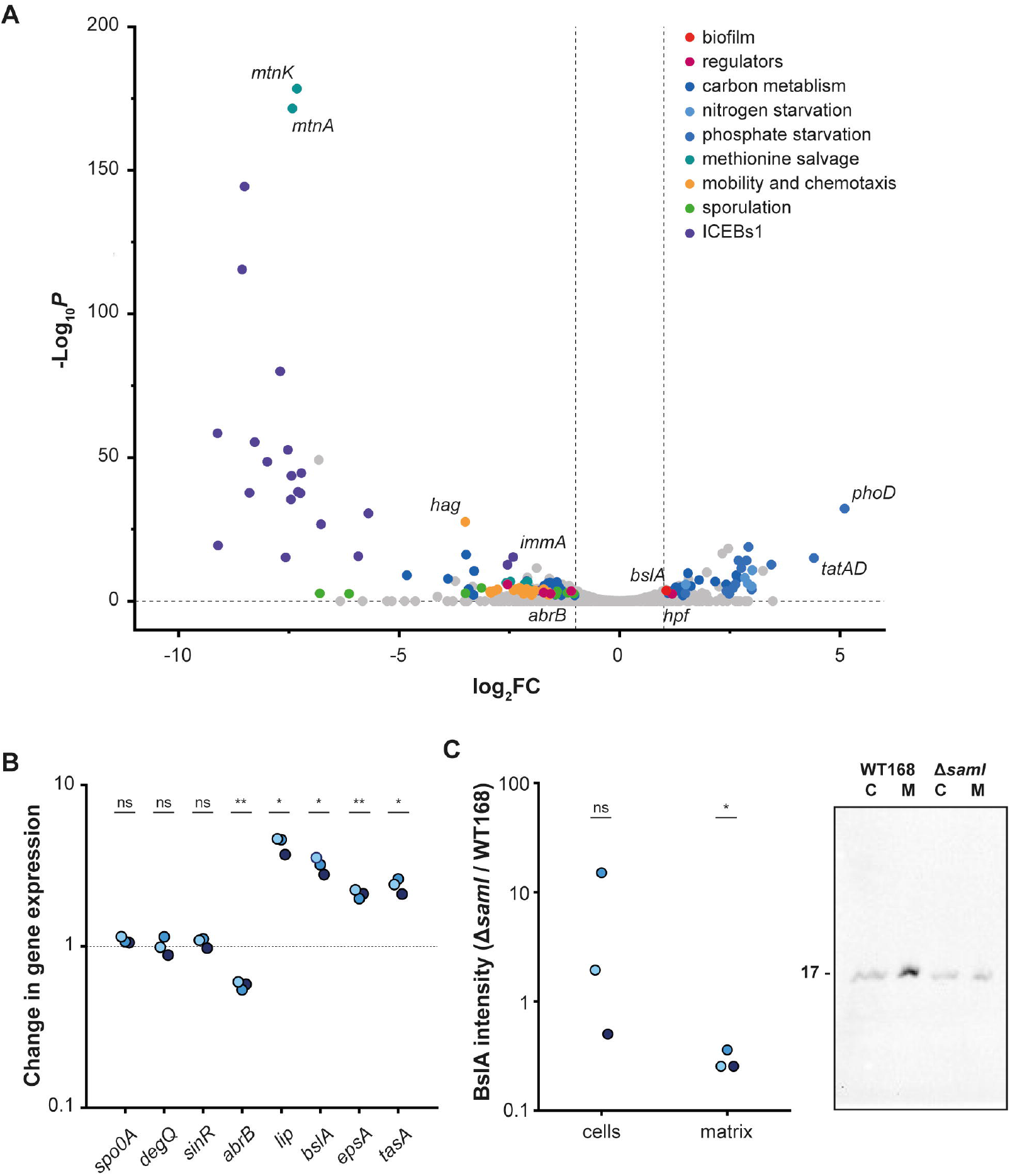
Transcriptional differences upon deletion of a *samI* mutant reveal changes in metabolism associated gene clusters and a reduction of BslA in the biofilm matrix. (A) Changes in gene expression upon *samI* deletion in a biofilm. RNAseq was performed with biofilms of WT168 and Δ*samI*. Significantly changed genes clusters are highlighted in different colors (see legend in upper right corner). (B) Changes in gene expression between WT168 and Δ*samI* in a biofilm measured with RT-qPCR. Expression levels of regulators of biofilm development and matrix genes were determined (and normalized to levels of *divIVA*). Three independent replicates are depicted in different colors. Significance was calculated using one-sample t-test, p-value ≤ 0.05. (C) Change in BslA levels between WT168 and Δ*samI* in a biofilm. BslA-His intensity (20.2 kDa) was determined within a biofilm separated into the cells (C) and the matrix (M) (see left graph). Exemplary Western blot is shown on the right. Three independent replicates are depicted in different colors. Significance was calculated using one-sample t-test, p-value ≤ 0.05.

Using reverse transcriptase qPCR, we confirmed that *bslA* transcription as well as the transcription of other matrix genes were upregulated (*epsA* and *tasA*), which is in line with the reduced expression of *abrB*, the negative regulator of biofilm genes and other genes including the *lip* gene, encoding an extracellular lipase (Figure 3B). Expression of other biofilm regulators (*spo0A, sinR*, and *degQ*) remained unchanged. Overall, the transcriptome analysis indicates that loss of SamI triggers a nutrient-starvation-like response, altering metabolism and stress pathways while simultaneously increasing matrix gene transcription.

To investigate why biofilms are not formed properly in a *samI* mutant we investigated protein levels of BslA with a *bslA* poly-histidine fusion integrated into the native locus of *bslA*. We tested if biofilm development was influenced by the additional His-tag and confirmed that it had no effect on BslA’s ability to form a hydrophobic coat during biofilm development (Figure S6A). Western blotting revealed that total BslA-His levels were on average decreased in Δ*samI* compared to WT168 (Figure S6B). Further separation into a fraction containing the biofilm matrix and the cellular fraction revealed that BslA-His was always reduced in the matrix of Δ*samI*, while BslA-His levels within the cell showed strong variations (Figure 3C). Thus, higher *bslA* transcription did not consistently translate into efficient protein production and secretion giving rise to the observed biofilm phenotype of Δ*samI*. If general stress response upon starvation leads to a reduction of protein synthesis this should also apply to other, not related proteins. Therefore, we used a translational fluorescent fusion to observe protein expression in more detail. The polar scaffold protein DivIVA is constitutively expressed in *B. subtilis* (Trip *et al*., 2013). The translational *divIVA* fusion exhibit a drastically reduced fluorescent intensity in a Δ*samI* background, indicating that protein expression is generally reduced in this strain background (Figure S7).

In total, Δ*samI* leads to a decrease of transcription of *abrB* in biofilms, and thus an increase in matrix gene transcription in biofilms. At the same time, a starvation-induced phenotype and a higher transcription of the hibernation factor *hpf* is observed in Δ*samI* that leads to reduced protein synthesis, and as a result decreased amounts of BslA are found in the matrix. Together, the transcriptome pattern indicates that loss of SamI induces a global nutrient-stress response, which decreases translational capacity. This stress response explains why matrix gene transcripts increase (due to reduced *abrB*), while the actual accumulation of BslA protein in the matrix is nevertheless reduced and in turn does not support correct biofilm maturation.

### SamI impacts motility

During the transcriptional analysis, a reduction in motility-associated genes was observed as well (Figure 3). Therefore, we wanted to validate a potential implication on bacterial motility experimentally. Using a fluorescent fusion of FliM-GFP, the number of flagella basal bodies in WT168 and Δ*samI* biofilms was determined (Figure 4A and B). In both the center and the periphery of biofilms, a reduced number of FliM-GFP foci was detected in Δ*samI*. While nearly 80 % of WT168 cells contained five FliM-GFP foci or more, only 30 % of Δ*samI* cells contained that many foci in cells from the periphery of the biofilm. The number of cells with no foci more than doubled, from 14 % to 39 %, in the Δ*samI* mutant. In the center of the biofilm, more than 80 % of Δ*samI* cells contained no foci, and only a minor number of cells contained one or more foci. In WT168, approximately 25 % of cells contained no FliM-GFP foci. The remaining cells all contained a smaller number of foci, with the highest fraction having one focus (24 %), followed by two foci (21 %), and three foci (13 %).

**Figure 4:**
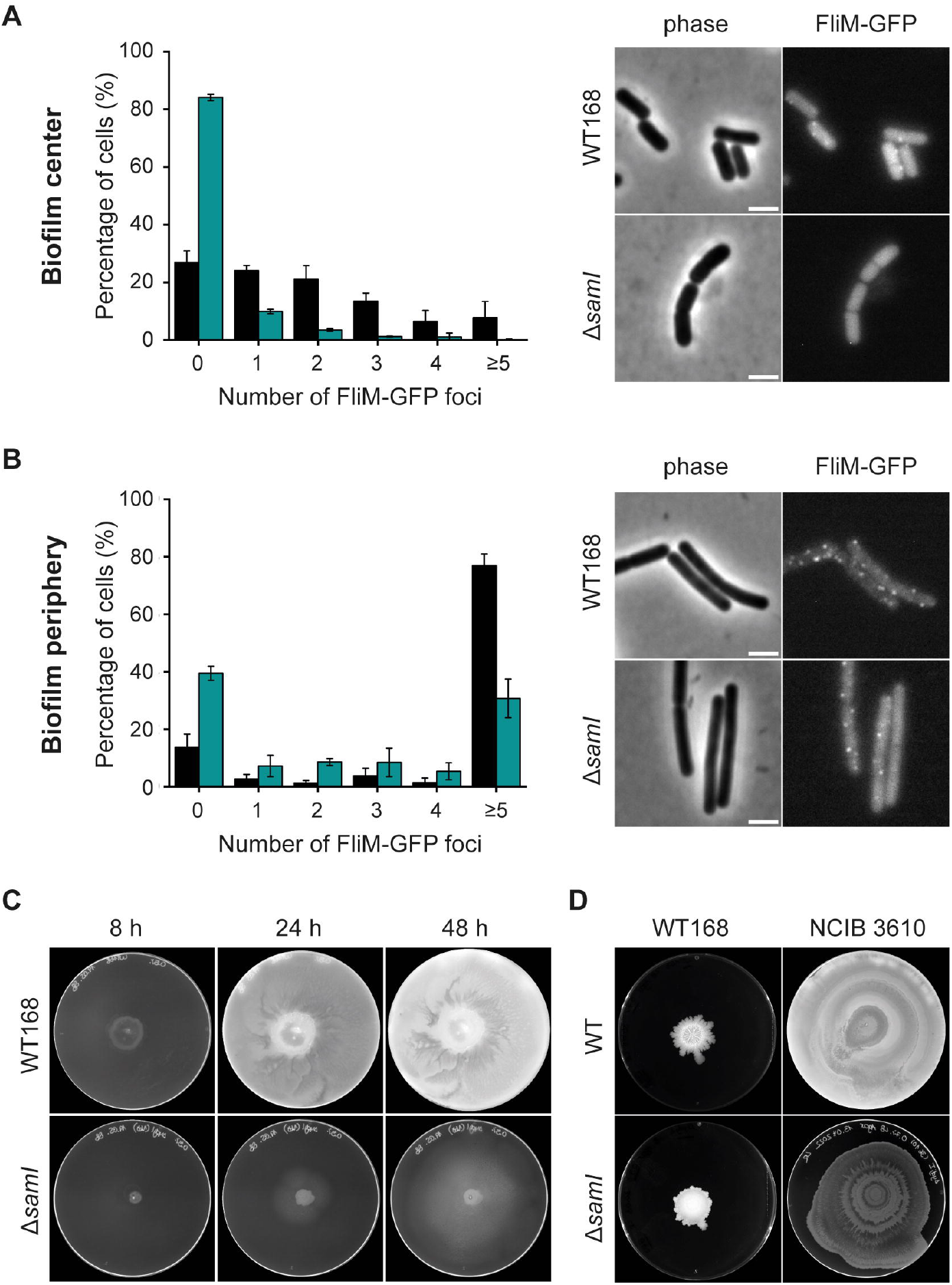
Loss of *samI* results in reduced motility. Number of FliM-GFP foci in WT168 (black) and Δ*samI* (teal) were determined in central (A) and peripheral (B) areas of biofilms after 48 h (average and standard derivation of three independent replicates, n = 618-1285 cells). Images of exemplary cell are shown on the right (scale: 2 µm) (C) Swimming progress of WT168 and Δ*samI* is shown after 8, 24, and 48 h for one exemplary replicate (N = 3, exemplary assay is shown, petri dish diameter: 92 mm). (D) Swarming progress after 48 h of wild type and Δ*samI* in WT168 and NCIB 3610 backgrounds is shown for one exemplary replicate (N = 3, exemplary assay is shown, petri dish diameter: 92 mm).

Therefore, we investigated SamI’s influence on motility further. *B. subtilis* has different modes of motility depending on the physical properties of its environment (Kearns & Losick, 2003, Kearns, 2010). In liquid environments, *B. subtilis* prefers to swim, a unicellular mode of motility exhibited by highly flagellated cells (Kearns, 2010). In conditions of higher viscosity, *B. subtilis* secretes surfactin, a lipopeptide that lowers the surface tension, and thereby facilitates movement of *B. subtilis* cell clusters (Kearns & Losick, 2003). This mode of motility is called swarming. During swimming assays, a reduced swimming ability of Δ*samI* was observed compared to WT168 (Figure 4C). For the swarming assays we additionally included the undomesticated wild type NCIB 3610, as the WT168 background used in this study up until now has a mutation in genes essential for surfactin production (Calvio *et al*., 2005, Zeigler *et al*., 2008, Patrick & Kearns, 2009). The strains with the WT168 background developed complex colonies during this assay due to their inability to swarm (Figure 4D). Here, a reduction of architectural complexity was observed as well for Δ*samI* similar to the reduced complexity during biofilm development observed before. Strains with the NCIB 3610 background, revealed that deletion of *samI* led to a reduction of swarming (Figure 4D).

In summary, deletion of *samI* greatly reduces motility in *B. subtilis* in swimming and swarming conditions in the laboratory background WT168 as well as the natural wild type NCIB 3610. In line with this phenotype, a reduction in transcript levels of the motility and chemotaxis regulon was observed in Δ*samI* biofilms, and a reduced number of basal bodies was detected during biofilm development. Together, these results indicate that SamI is required for both the proper transcription of motility genes and the formation of flagellar structures.

## 4. Discussion

*B. subtilis* adapts to fluctuating environments by differentiating into distinct cell types capable of coping with specific stresses (Vlamakis *et al*., 2008, Lopez & Kolter, 2010, Qin *et al*., 2022). During exponential growth, a subset of cells adopts a motile phenotype, enabling dispersal into new niches (Kearns & Losick, 2005). As cell density increases and nutrients become limiting, cells may differentiate into competent cells, cannibal cells, spores, or matrix-producing cells that form protective biofilms (Lopez *et al*., 2009, Mirouze & Dubnau, 2013). Mature biofilms are heterogeneous, comprising of motile, matrix-producing, and sporulating subpopulations that collectively shape biofilm architecture (Vlamakis *et al*., 2008).

Here, we show that deletion of *samI*, which encodes an SPFH domain protein within the σ^W^-regulated *pspA-samGHI* operon, leads to systemic consequences within a biofilm resulting in incomplete biofilm development: Deletion of *samI* produces smooth, non-structured colonies. This phenotype is not observed upon single deletion of *pspA, samG, samH*, or the entire operon (Figures 1 and 2). Notably, co-deletion of *samH* suppresses the Δ*samI* phenotype, and partial complementation of Δ*samI* requires deletion of *samG* (Figures 1 and S3). These observations indicate that the defect in the hydrophobic biofilm formation does not result from simple loss of SamI function, but rather from unrestrained signaling by the SamH, which is further modulated by the presence of SamG. Thus, in wild type cells, SamI restrains SamGH activity, preventing chronic envelope-stress signaling (Figure 5). When SamI is absent, SamGH presumably becomes constitutively active, inducing a persistent stress response that drives metabolic rewiring, including changes to nutrient-starvation programs.

**Figure 5:**
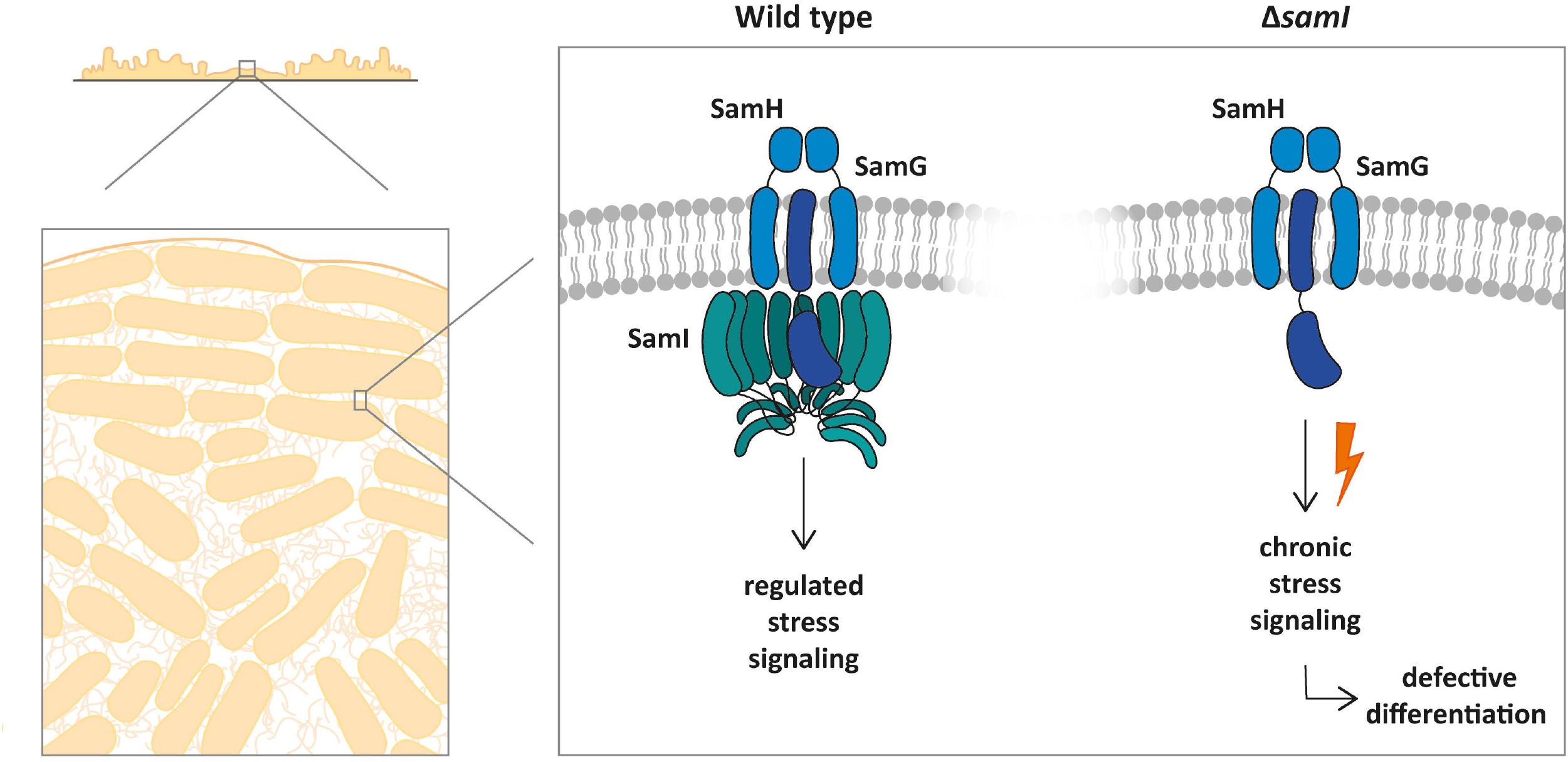
Schematic model of SamI’s impact on biofilm development. In the wild type (left) SamI regulates the membrane complex SamGH resulting in a normal stress response during biofilm development. In absence of *samI* (right) the SamGH complex is constitutively active leading to a persistent stress response resulting in metabolic changes that prevent BslA layer formation and flagellar basal body assembly. Oligomerization states are based on structure predictions from AlphaFold 3 predicting a monomer for SamG and a dimer for SamH. Predictions of SamI and unpublished data indicated a higher ordered oligomer.

This regulatory logic parallels the phage shock protein (Psp) system in *E. coli* (Kleerebezem *et al*., 1996, Jovanovic *et al*., 1996, Dworkin *et al*., 2000, Flores-Kim & J., 2016, Popp *et al*., 2022). In that system, PspB and PspC form a membrane-associated sensor, while PspA acts as a negative regulator that restrains the transcription factor PspF. Loss of PspA leads to constitutive stress signaling, mimicking the effect of chronic SamGH activation in Δ*samI*. Although SamI and PspA differ structurally, a SPFH domain scaffold versus coiled-coil regulator, the organizational principle is conserved: a membrane sensor complex is held in check by a regulatory subunit to prevent persistent stress responses. The *E. coli* PspA is, however, not only a regulator of the phage shock system, but also the executing module. PspA is a bacterial ESCRT-III homolog that is likely involved in membrane dynamics (Kobayashi *et al*., 2007, Engl *et al*., 2009, Gupta *et al*., 2021, Junglas *et al*., 2021, Liu *et al*., 2021, Hudina *et al*., 2025, Junglas *et al*., 2025a, Junglas *et al*., 2025b). Similarly, the *B. subtilis pspA-samGHI* operon also contains a *pspA* homolog. Thus, in the firmicute system the regulation of the sensory module and the effector complex have been separated into two proteins, PspA and SamI, respectively.

Members of the SPFH domain family have been shown to frequently function as scaffolds that spatially organize and regulate membrane-associated complexes (Bramkamp & Lopez, 2015, Willdigg & Helmann, 2021, Bramkamp & Scheffers, 2023). Examples include HflK/C regulating the FtsH protease in *E. coli*, the SPFH-NfeD complex QmcA-YbbJ in *E. coli*, and the stomatin-like protein StlP controlling cellulose synthase complexes in *Streptomyces coelicolor* (Ma *et al*., 2022, Qiao *et al*., 2022, Tan *et al*., 2024, Zhong *et al*., 2025). SamI likely acts in a similar manner, modulating SamGH signaling and preventing chronic activation during multicellular development.

Further conceptual parallels exist between SamI/SamGH and the LiaRS two-component system in *B. subtilis*, which responds to antibiotic– and cell envelope–induced stress (Radeck *et al*., 2017). In the Lia system, LiaF inhibits the sensor kinase LiaS under non-stress conditions leaving LiaS phosphatase activity intact, preventing constitutive activation of the LiaRS regulon (Jordan *et al*., 2006). Only upon envelope perturbation is this inhibition relieved, triggering an adaptive stress response. This mirrors the function of SamI, which restrains SamGH signaling under normal conditions. Loss of SamI produces unrestrained SamGH activity, analogous to removal of LiaF, resulting in chronic signaling and maladaptive cellular responses. Together, these examples illustrate a conserved regulatory principle in *B. subtilis*: membrane-associated sensor complexes require inhibitory “brake” proteins to prevent constant stress signaling, whether in the context of biofilm development (SamI/SamGH) or antibiotic-induced envelope stress (LiaF/LiaRS).

Importantly, the regulation of membrane spanning signaling proteins mediated by SPFH domain protein is conserved also in eukaryotes. Flotillins and stomatins have been implicated in organizing membrane microdomains in eukaryotic cells (Langhorst *et al*., 2005, Browman *et al*., 2007, Lapatsina *et al*., 2012, Yokoyama & Matsui, 2020). Flotillin-1 and -2 regulate the levels of CD14 on macrophages, which recognizes LPS thus activating several TLR signaling pathways, including TLR4, of the innate immune response (Matveichuk *et al*., 2024). Also, Flotillin-1 positively regulates endosomal TLR3 signaling in endothelial cells by promoting ligand transport and caveolae-dependent receptor activation, thereby enabling inflammatory responses (Fork *et al*., 2014). These examples reveal the universal role of SPFH domain proteins in signaling pathways in pro- and eukaryotes.

Loss of SamI in *B. subtilis* triggers profound physiological consequences. Δ*samI* biofilms exhibit reduced translation, leading to a drastically reduced secretion of the matrix protein BslA and loss of biofilm architecture (Figures 1–3, S5–S6). Another consequence of the starvation phenotype is the downregulation of motility associated genes. Consequently, FliM-GFP basal bodies were found to be reduced, and both swimming and swarming were impaired in laboratory and undomesticated strains (Figure 4).

Further evidence for a starvation phenotype that affects cellular energy levels is the downregulation of ICEBs1 genes in Δ*samI* biofilms, indicating that horizontal gene transfer is suppressed under these conditions (Figure 3). This is likely a consequence of the starvation-like metabolic state observed in Δ*samI* cells. Maintenance of ICEBs1 within a population is an energy-intensive process requiring integration, excision, replication, and translocation of ICEBs1 (Auchtung *et al*., 2016). While replication can include multiple rounds of replication per donor, translocation is facilitated by a Type 4 Secretion System (T4SS) including the putative ATPases ConQ and ConE, responsible for coupling of the ICEBs1 to the T4SS and DNA transfer, respectively (Lee *et al*., 2010, Berkmen *et al*., 2010, Auchtung *et al*., 2016, Murthy *et al*., 2023). Under nutrient-limited or low-energy conditions, cells likely prioritize essential survival functions over horizontal gene transfer, leading to reduced expression of ICEBs1 genes. This finding highlights how the metabolic consequences of chronic SamGH signaling in the absence of SamI can directly impact cellular programs beyond biofilm formation, including adaptive gene exchange.

In conclusion, SamI functions as a regulator that restrains SamGH-mediated stress signaling during biofilm development. By preventing unrestrained activity, SamI ensures correct cellular responses during growth on surfaces (Figure 5). This is important at the population level to allow the fine-tuned differentiation of different developmental pathways. In turn, chronic stress induced by unregulated signal transduction leads to misregulation of the signal cascades governing these programs. The analogy with the *E. coli* Psp system highlights a conserved design principle in bacteria: sensor complexes require regulatory subunits to avoid maladaptive chronic stress signaling. In *B. subtilis*, SamI exemplifies this regulatory principle, enabling cells to integrate stress perception with multicellular development.

## 6. Acknowledgments

This project was funded by the DFG (BR2915/7-1). The SEM microscope was purchased through grant INST 257/688-1 (FUGG). We thank Anika Lassota for the construction of the plasmids pKill*-samI-halo-spec* and pKill*-spec-halo-samI* and we are grateful to Ulrike Voigt and Tanja Zabel (Central Microscopy, CAU) for their technical assistance with sample preparation for electron microscopy.

We thank the NGS team of the Bielefeld University Omics CF NGS Unit (in development) and the technical staff of the CeBiTec Technology Platform Genomics, in particularly Eva Schulte-Bernd, Yvonne Kutter, and Katharina Hanuschka, for their technical assistance.

Additionally, we would like to thank Oscar P. Kuipers for the kind gift of the plasmid pDRyuaB2. We thank Daniel B. Kearns for the kind gift of the strain DS8521, Jeffrey Errington for the kind gift of strain 1803, and Thorsten Mascher for the kind gift of the *B. subtilis* strain Δ*pspA-samGHI*.

