## Supplemental Material for "An SPFH Protein Couples Membrane Stress to Differentiation in *Bacillus subtilis*"

### 1. Supplementary figures

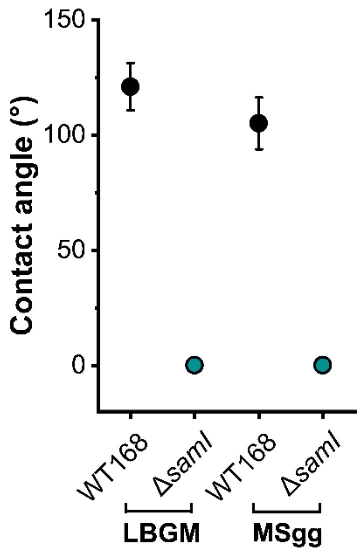


**Supplementary Figure S1: Deficient biofilm phenotype of Δ*samI* is independent of medium.** Surface hydrophobicity of Δ*samI* biofilms is deficient on full (LBGM) and minimal medium (MSgg). Average contact angle and standard derivation was determined in three independent replicates.

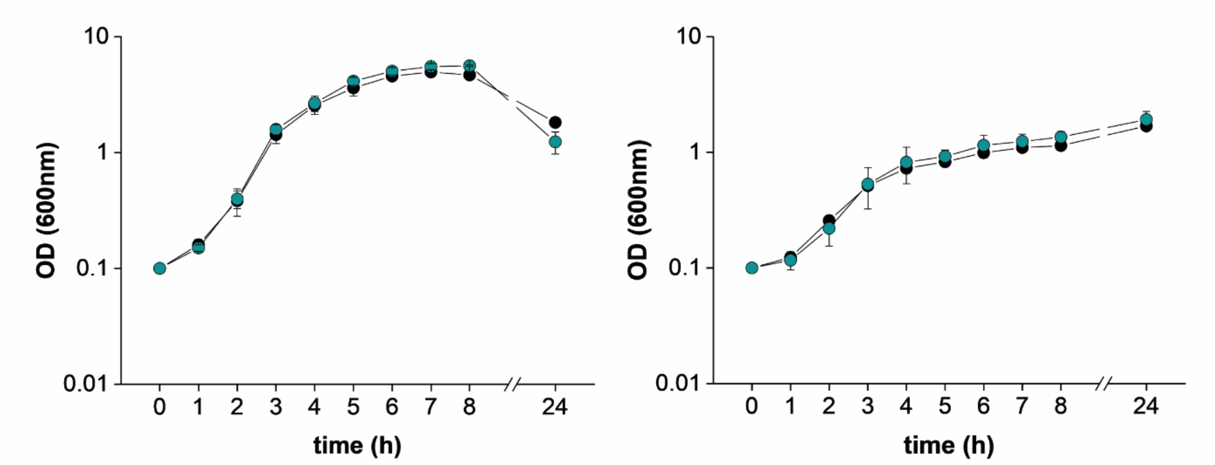


**Supplementary Figure S2: *SamI* mutant grows indistinguishable from WT168 in liquid cultures.** Growth curves of WT168 and Δ*samI* in full medium (right) and minimal medium (left) over a time course of 24 h. Average and standard derivation are depicted of three independent replicates.

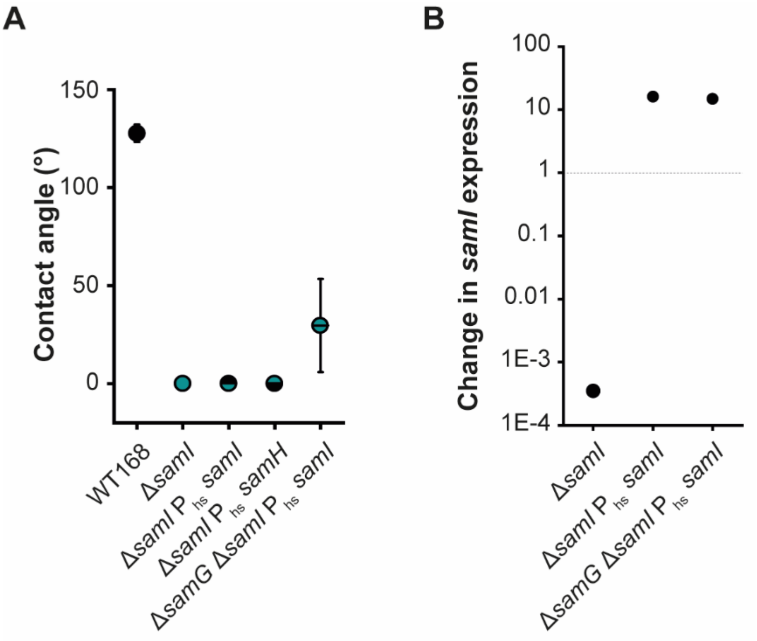


**Supplementary Figure S3: Absence of *samG* leads to partial complementation of *samI*.** (A) Surface hydrophobicity in Δ*samI* can only be restored in complementation strains when SamI membrane anchor, SamG, is missing. Contact angle and standard derivation was determined of three independent replicates of WT168, Δ*samI*, and its complementation strains. (B) Transcript level of *samI* in the corresponding strains normalized to level of *samI* in WT168.

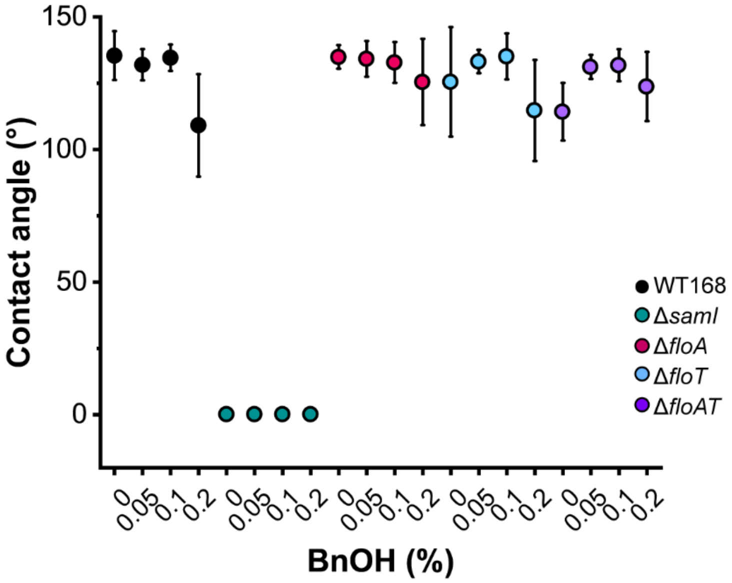


**Supplementary Figure S4: Membrane fluidization does not improve Δ*samI's* biofilm phenotype.** Surface hydrophobicity of SPFH mutants of LBGM supplemented with 0 % to 0.2 % benzyl alcohol. Contact angle and standard derivation was determined of three independent replicates.

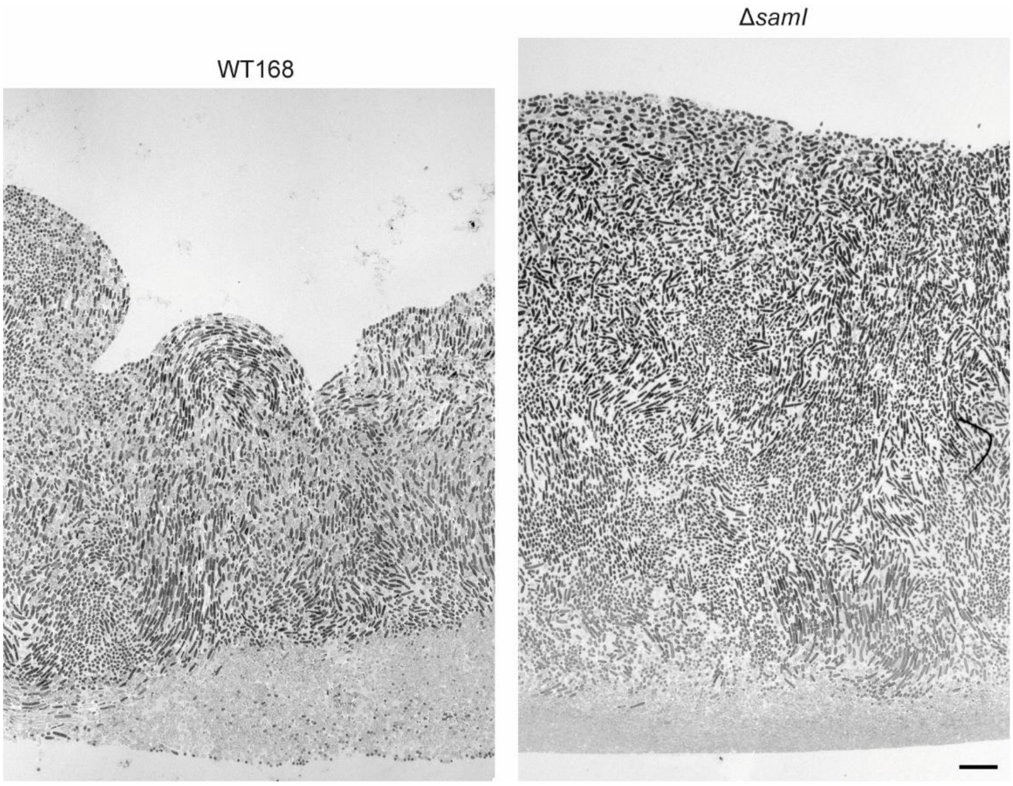

**Supplementary Figure S5: Biofilms of WT168 and Δ*samI* have a layer of empty cells at the bottom of the biofilm.** TEM images of showing an overview over cross section of WT168 (left) and Δ*samI* (right) biofilms. On the bottom of the biofilms a layer of empty cell envelopes is visible (scale: 10 µm).

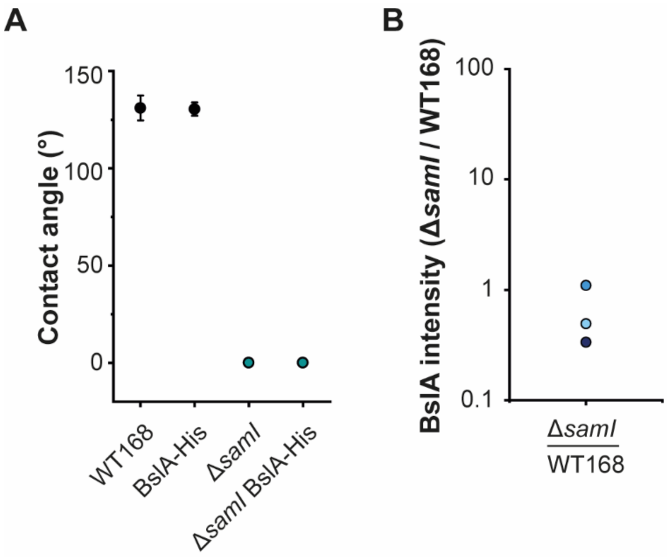


**Supplementary Figure S6: BslA-His does not influence biofilm hydrophobicity and Δ*samI* exhibit reduced BslA-His levels.** (A) Surface hydrophobicity of WT168 and Δ*samI* is not impacted by His tag on BslA. Contact angle and standard derivation was determined of three independent replicates. (B) Change in BslA-His level between WT168 and Δ*samI* isolated from biofilms. Three independent replicates are depicted in different colors.

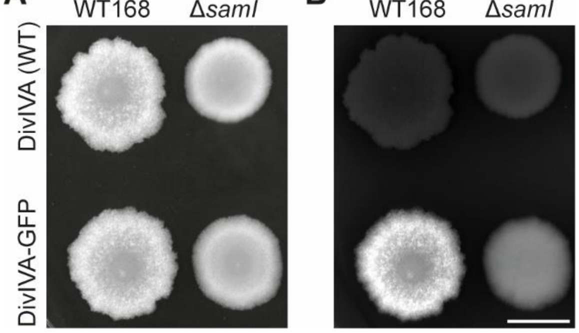

**Supplementary Figure S7: DivIVA-GFP levels are reduced in a Δ*samI* biofilm.** *divIVA-GFP* intensity in biofilms of WT168 and Δ*samI* background strains (scale: 1 cm). Biofilm morphology is shown in (A) and GFP intensity in (B).

#### 2. Supplementary tables

**Table S1:** *Bacillus subtilis* strains used in this study.

| Strain number | Name | Genotype/description | Reference |
|---|---|---|---|
| SBB001 | WT168 | *trpC2* | Laboratory collection |
| SBB002 | WT168 Δ*floT* | *trpC2 yuaG::pMUTIN4 erm* | This work |
| SBB003 | WT168 Δ*samI* | *trpC2 samI::kan* | This work |
| SBB004 | WT168 Δ*pspA* | *trpC2 pspA::kan* | This work |
| SBB005 | WT168 Δ*samH* | *trpC2 samH::kan* | This work |
| SBB006 | WT168 Δ*samG* | *trpC2 samG::kan* | This work |
| SBB007 | WT168 Δ*pspA-samGHI* | *trpC2, pspA-samGHI::(*$P_{pspA}$ *erm)* | This work |
| SBB008 | DSM10 / NCIB 3610 | Undomesticated wild type strain | Laboratory collection |
| SBB009 | NCIB 3610 Δ*samI* | *samI::kan* | This work |
| SBB041 | WT168 Δ*ytrF* | *trpC2 ytrF::kan* | This work |
| SBB060 | WT168 Δ*samI* $P_{hs}$-*samI* | *samI::kan amyE::*$P_{hyperspank}$*-samI spec* | This work |
| SBB067 | WT168 Δ*samG* Δ*samI* $P_{hs}$-*samI* | *samG::erm samI::kan amyE::*$P_{hyperspank}$*-samI spec* | This work |
| SBB070 | WT168 Δ*samG* | *samG::erm* | This work |
| SBB071 | WT168 Δ*floA* | *trpC2 yqfA::tet* | This work |
| SBB094 | WT168 Δ*samI* | Δ*samI* | This work |
| SBB100 | WT168 Δ*floT* Δ*floA* | *trpC2 yuaG::pMUTIN4 erm yqfA::tet* | This work |
| SBB127 | WT168 BslA-His | *trpC2 bslA::bslA-6xHis-cat* | This work |
| SBB129 | WT168 Δ*samI* BslA-His | *trpC2 samI::kan bslA::bslA-6xHis-cat* | This work |
| SBB131 | WT168 Δ*ytrF* BslA-His | *trpC2 ytrF::kan bslA::bslA-6xHis-cat* | This work |
| SBB133 | WT168 DivIVA-GFP | *trpC2 divIVA::divIVA-GFP cat* | This work |
| SBB134 | WT168 Δ*samI* DivIVA-GFP | *trpC2 samI::kan divIVA::divIVA-GFP cat* | This work |
| SBB136 | WT168 Δ*samI* Δ*pspA* | trpC2 Δ*samI pspA::kan* | This work |
| SBB137 | WT168 Δ*samI* Δ*samG* | trpC2 Δ*samI samG::kan* | This work |
| SBB138 | WT168 Δ*samHI* | *trpC2 samH-samI::erm* | This work |
| DS8521 | WT168 Δ*fliM,* $P_{fla/che}$ FliM-GFP | Δ*fliM amyE::*$P_{fla/che}$*-fliM–GFP spec* | This work |
| SBB149 | WT168 Δ*fliM* | *trpC2 fliM::kanR* | This work |
| SBB150 | WT168 Δ*samI* Δ*fliM* | *trpC2* Δ*samI fliM::kanR* | This work |
| SBB151 | WT168 Δ*fliM* $P_{fla/che}$ FliM-GFP | *trpC2 fliM::kanR amyE::*$P_{fla/che}$*-fliM–GFP spec* | This work |

| | | | |
|---|---|---|---|
| SBB152 | WT168 Δ*samI* Δ*fliM* $P_{fla/che}$ FliM-GFP | *trpC2* Δ*samI fliM::kanR amyE::*$P_{fla/che}$*-fliM–GFP spec* | This work |
| SBB159 | WT168 Δ*samGH* | *trpC2 samG-samH::erm* | This work |
| SBB160 | WT168 Δ*samGHI* | *trpC2 samG-samH-samI::erm* | This work |
| SBB193 | WT168 Δ*samI* $P_{hs}$-*samH* | *trpC2 samI::kan amyE::*$P_{hyperspank}$*-samH spec* | This work |
| DB003 | WT168 Δ*floT* | *trpC2 yuaG::pMUTIN4 erm* | (Donovan and Bramkamp 2009) |
| BB001 | WT168 Δ*floA* | *trpC2 yqfA::tet* | (Bach and Bramkamp 2013) |
| 1803 | DivIVA-GFP | *divIVA::(*$P_{divIVA\text{-}gfp}$ *divIVA*$^{+}$ *cat)* | (Edwards, Thomaides, and Errington 2000) |
| DS8521 | WT168 Δ*fliM,* $P_{fla/che}$ FliM-GFP | Δ*fliM amyE::*$P_{fla/che}$*-fliM–GFP spec* | (Guttenplan, Shaw, and Kearns 2013) |
| | WT168 Δ*pspA-samGHI* | *trpC2, pspA-samGHI::(*$P_{pspA}$ *erm)* | Gift from T. Mascher |

**Table S2:** *Escherichia coli* strains used in this study.

| Strain number | Name | Reference |
|---|---|---|
| SBE002 | DH5α | Laboratory collection |
| SBE005 | DH5α pKill | Laboratory collection |
| SBE011 | DH5α pDR111-*samI* | This work |
| SBE013 | DH5α pDR244 | (Koo et al. 2017) |
| SBE023 | DH5α pKill-*samI-halo-spec* | A. Lassota |
| SBE024 | DH5α pKill-*spec-halo-samI* | A. Lassota |
| SBE029 | DH5α pKill-Δ*samHI* | This work |
| SBE030 | DH5α pKill-Δ*samGH* | This work |
| SBE031 | DH5α pKill-Δ*samGHI* | This work |
| SBE041 | DH5α pDR111-*samH* | This work |
| | XL Blue1 pDRyuaB2 | (Kovacs and Kuipers 2011) |

**Table S3:** Plasmids used in this study.

| Name | Description | References |
|---|---|---|
| pKill | pUC18mut with *txpA* (*yqdB*) gene | Laboratory collection |
| pDR244 | temperature-sensitive plasmid with constitutively expressed Cre recombinase | (Koo et al. 2017) |
| pDR111-*samI* | For ectopic expression of *samI* under the $P_{hyperspank}$ promoter (IPTG inducible) | This work |
| pKill-Δ*samHI* | For generation of a double deletion of *samH* and *samI* | This work |
| pKill-Δ*samGH* | For generation of a double deletion of *samG* and *samH* | This work |
| pKill-Δ*samGHI* | For generation of a triple deletion of *samG*, *samH*, and *samI* | This work |
| pDR111-*samH* | For ectopic expression of *samH* under the $P_{hyperspank}$ promoter (IPTG inducible) | This work |
| pKill-*samI-halo-spec* | C-terminal *samI-halo* fusion | A. Lassota |
| pKill-*spec-halo-samI* | N-terminal *samI-halo* fusion | A. Lassota |

| | | |
|---|---|---|
| pDRyuaB2 | For ectopic expression of *bslA* under the $P_{hyperspank}$ promoter (IPTG inducible) | (Kovacs and Kuipers 2011) |

**Table S4:** Oligonucleotides used in this study.

| Primer Number | Primer | Sequence 5'→ 3' |
|---|---|---|
| SB001 | samI_500bp_upstream_fw | ACGACCAAAAGCACAAAC |
| SB027 | pDR111_fwd | AAGCTTAATTGTTATCCGCT |
| SB028 | pDR111_rev | GCATGCAAGCTAATTCGGTG |
| SB029 | samI_pDR111_fwd | CACCGAATTAGCTTGCATGCTTATACAAGCTTCTGGCC |
| SB030 | samI_pDR111_rev | AGCGGATAACAATTAAGCTTAAAGGGAGAGGCGTAATG |
| SB035 | pDR111_seq_fw | GGCAAGAACGTTGCTCGAG |
| SB036 | pDR111_seq_rv | TCAGCCGACTCAAACATCAA |
| SB037 | pKill_fwd | GCTTGGCACTGGCCGTCGTTT |
| SB038 | pKill_rev | TTGCATGCCTGCAGGTCGACT |
| SB052 | samI_500bp_downstream_rv | CCCCATCCTTCATTTTTACCA |
| SB080 | divIVA_fw | GAGCTTGAAGCGAAAGTCAATGAG |
| SB081 | divIVA_rv | TCGTTGATAATGCGATCAGCG |
| SB086 | spo0A_fw | TGGGCAGGAAGATGTCACG |
| SB087 | spo0A_rv | CACGATGCGTCACACTGCTG |
| SB088 | abrB_fw | CCTATCGAACTGCGTCGTACTC |
| SB089 | abrB_rv | CCGCCTGCAAGTTTAAGGTTATC |
| SB090 | lip_fw | CAATCCAGTCGTTATGGTTCAC |
| SB091 | lip_rv | CGTGATAATACCGGTCCATTG |
| SB092 | degQ_fw | CAATTGTTATTCCGACTCGAACTTG |
| SB093 | degQ_rv | AATTGTATTTATCGAGTTGATCAATGC |
| SB098 | tasA_fw | CCGGGAGATAAGTTGACAAAGG |
| SB099 | tasA_rv | GTAGCCATTGCCGCCCTCTT |
| SB100 | bslA_fw2 | GTTACTCGTTTCTGCACC |
| SB101 | bslA_rv2 | GGACGGTAAGTCAATTCG |
| SB102 | sinR_fw | GTCTCCGCTGTTCTGGACG |
| SB103 | sinR_rv | CGGATGTCATCGCATCGC |
| SB104 | bslA_fwd | CCTCTAGAGTCGACCTGCAGGCATGCAAATGAAACGCAAATTATTATCTTCTTTGGC |
| SB105 | bslA_GG_6xHis_rev | ATTTTTATCTAAAGATGATGATGATGATGATGCCCTCCGTTGCAACCGCAAGGCTGAG |
| SB106 | catR_GG_6xHis_fwd | CTTGCGGTTGCAACGGAGGGCATCATCATCATCATCATCTTTAGATAAAAATTTAGGAGGCATATC |
| SB111 | catR_rev | ACCGGTCTTTTACATTATAAAAGCCAGTCATTAGGC |
| SB114 | bslA_dw_fwd_2 | ATAATGTAAAAGACCGGTTAACGCC |
| SB115 | bslA_dw_rev_2 | GTTGTAAAACGACGGCCAGTGCCAAGCCGATGGCAACAATCACATATTC |
| SB116 | samI_fw | GCCAAGGGAATTCAAGAGGACTTA |
| SB117 | samI_rv | CCAACCATACCGTAGGAAGCG |
| SB118 | pKill_seq_fw | GGATCCTCTAGAGTCGACC |
| SB119 | pKill_seq_rv | CTGCAAGGCGATTAAGTTG |
| SB137 | up_ldh | TACCCTGGAAGGATGATTA |
| SB138 | dw_ldh | TACTCTAAAGTTGCGGTTAG |

| | | |
|---|---|---|
| SB160 | samI-pKill_GH(I)_rv | ATAAAGCATTTTGCATGCCTGCAGGTCGAC |
| SB161 | pspA_GH(I)_fw | AGGCATGCAAAATGCTTTATTGGACAAGGC |
| SB162 | pspA_GH(I)_rv | TCTCGCCTGCTTACTTATCGAGCATCATTTTCG |
| SB163 | ermR_GH(I)_fw | CGATAAGTAAGCAGGCGAGAAAGGAGAGAG |
| SB164 | ermR_GH_rv | AAAACGACATCGAGGCTCCTGTCACTGCTTC |
| SB165 | samI-dw-pKill_(G)HI_fw | GAATCATACAGTGAAAAGTCCGGAG |
| SB166 | ermR_(G)HI_rv | GACTTTTCACTGTATGATTCCGAGGCTCCTGTCACTGCTTC |
| SB167 | samI-dw-pKill_HI_rv | CAGTTTGCTCTTGCATGCCTGCAGGTCGAC |
| SB168 | samG_HI_fw | AGGCATGCAAGAGCAAACTGTACCAGAGTCC |
| SB169 | samG_HI_rv | TCTCGCCTGCTCAAAATCCGCCTCCCATCATC |
| SB170 | ermR_HI_fw | CGGATTTTGAGCAGGCGAGAAAGGAGAGAG |
| SB257 | pDR111_rev | TACGCCTCTCCCTTTAAGCTTA |
| SB258 | samH_fwd | AGCTTAAAGGGAGAGGCGTAATGCGTGGATTTTTTGGG |
| SB259 | samH/GH_rev | CACCGAATTAGCTTGCATGCTTAAAAACTGCCCCGGCTC |

**Table S5:** Average contact angles and corresponding standard derivation (SD).

| Strain | Average ± SD |
|---|---|
| Figure 1B | |
| WT168 | 125.7° ± 6.58° |
| Δ*pspA* | 130.4° ± 12.27° |
| Δ*samG* | 130.2° ± 7.30° |
| Δ*samH* | 131.5° ± 8.03° |
| Δ*samI* | 0.0° ± 0.00° |
| Δ*pspA* Δ*samI* | 5.7° ± 11.17° |
| Δ*samG* Δ*samI* | 0.0° ± 0.00° |
| Δ*samH* Δ*samI* | 116.8° ± 10.45° |
| Δ*samG* Δ*samH* | 119.5° ± 5.00° |
| Δ*samG* Δ*samH* Δ*samI* | 119.5° ± 5.27° |
| Δ*pspA-samGHI* | 131.0° ± 9.28° |

| Figure S2 | |
|---|---|
| WT168, LBGM | 120.9° ± 10.24° |
| Δ*samI*, LBGM | 0.0° ± 0.00° |
| WT168, MSgg | 105.0° ± 11.23° |
| Δ*samI*, MSgg | 0.0° ± 0.00° |

| Figure S3A | |
|---|---|
| WT168 | 127.8° ± 4.44° |
| Δ*samI* | 0.0° ± 0.00° |
| Δ*samI* $P_{hyper}$ *samH* | 0.0° ± 0.00° |
| Δ*samI* $P_{hyper}$ *samI* | 0.0° ± 0.00° |
| Δ*samG* Δ*samI* $P_{hyper}$ *samI* | 29.7° ± 23.80° |

| Figure S4 | |
|---|---|
| WT168, 0 % BnOH | 135.4° ± 9.22° |
| WT168, 0.05 % BnOH | 132.0° ± 5.89° |
| WT168, 0.1 % BnOH | 134.6° ± 5.00° |
| WT168, 0.2 % BnOH | 109.1° ± 19.31° |

| | |
|---|---|
| Δ*samI*, 0 % BnOH | 0.0° ± 0.00° |
| Δ*samI*, 0.05 % BnOH | 0.0° ± 0.00° |
| Δ*samI*, 0.1 % BnOH | 0.0° ± 0.00° |
| Δ*samI*, 0.2 % BnOH | 0.0° ± 0.00° |
| Δ*floA*, 0 % BnOH | 134.9° ± 4.46° |
| Δ*floA*, 0.05 % BnOH | 134.2° ± 6.73° |
| Δ*floA*, 0.1 % BnOH | 132.8° ± 7.70° |
| Δ*floA*, 0.2 % BnOH | 125.5° ± 16.24° |
| Δ*floT*, 0 % BnOH | 125.5° ± 20.61° |
| Δ*floT*, 0.05 % BnOH | 133.2° ± 4.41° |
| Δ*floT*, 0.1 % BnOH | 135.1° ± 8.67° |
| Δ*floT*, 0.2 % BnOH | 114.8° ± 19.03° |
| Δ*floAT*, 0 % BnOH | 114.3° ± 10.85° |
| Δ*floAT*, 0.05 % BnOH | 131.2° ± 4.52° |
| Δ*floAT*, 0.1 % BnOH | 131.8° ± 6.02° |
| Δ*floAT*, 0.2 % BnOH | 123.8° ± 13.05° |

| Figure S5A | |
|---|---|
| WT168 | 131.1° ± 6.44° |
| WT168, BslA-His | 130.5° ± 3.42° |
| Δ*samI* | 0.0° ± 0.00° |
| Δ*samI*, BslA-His | 0.0° ± 0.00° |

**Table S6:** RNA-Sequencing analysis

This table is shown in a separate Excel file. The table shows RNAseq analysis with fold changes of transcripts comparing *samI* mutant biofilms to WT168 biofilms.
